# Host use preference of fruit flies (Diptera: Tephritidae) among selected cucurbitaceous vegetables in Morogoro, Eastern-Central Tanzania

**DOI:** 10.1101/2023.04.20.537659

**Authors:** Petronila Tarimo, Sija Kabota, Maulid Mwatawala, Ramadhan Majubwa, Abdul Kudra, Massimiliano Virgilio, Kurt Jordaens, Marc De Meyer

## Abstract

Fruit flies (Diptera, Tephritidae) represent a major threat to cucurbit production in Tanzania. They cause huge economic losses on cucurbit crops. Information on the infestation and yield loss caused by fruit flies on cucurbit crops is vital in designing a sound fruit fly management program. We identified the fruit fly species attacking cucurbit crops (Cucurbitales: Cucurbitaceae): cucumber (*Cucumis sativus* L.), watermelon (*Citrullus lanatus* [Thunb.] Matsum. & Nakai) and squash (*Cucurbita moschata* D.) in the Morogoro region and quantified their incidence and infestation per crop species. A weekly sampling of ten fruits per crop was repeated twice per plot for each zone for five consecutive weeks between March and October 2020. The number of fruit flies emerging from the collected fruits was quantified and infestation per crop species was determined. We observed significant differences in infestation between cucurbit species, across zones and seasons. Squash showed the highest infestation of *Zeugodacus cucurbitae* (Coquillet) in both zones followed by watermelon. Watermelon was highly infested by *Dacus vertebratus* (Bigot) in the plateau zone followed by squash. Squash showed the highest infestation of *Dacus ciliatus* Loew in the plateau zone followed by watermelon. Cucumber had the lowest infestation for all fruit fly species.

## 1. Introduction

Cucurbitaceous crops (Cucurbitales: Cucurbitaceae) are among the most important vegetable crops grown worldwide, of their nutritional and economic value. In sub-Saharan Africa, cucurbit crops are of considerable economic importance for local diet, employment, and income generation [11, 12]. The commonly grown cucurbit species in Africa include cucumber (*Cucumis sativus* L.), watermelon (*Citrullus lanatus* [Thunb.] Matsum. & Nakai) and squash (*Cucurbita moschata* D.) [11]. Despite their economic importance, the production of cucurbit crops in Africa faces a number of challenges, including insect pest infestation, diseases, and climate change.

Fruit flies (Diptera: Tephritidae) are economically important pests of cucurbit crops, causing yield losses, ranging from 30% to 100% in tropical and sub-tropical regions [1, 9,24]. Considerable efforts by both governmental bodies, research institutions, and individuals have been made to implement programs including studies to manage and monitor the spread and impact of these pests in Africa [6,7,13-15]. Despite such efforts, losses due to fruit fly infestations continue to be felt with severe impacts and consequences being experienced much more in developing countries due to inadequate control of fruit flies.

In Africa, knowledge related to the host range of cucurbit-infesting fruit fly species throughout their geographic ranges is generally limited to a few species. Most studies on host range and preference focused on the invasive *Zeugodacus cucurbitae* (Coquillet) [12,13, 15] Only a few studies (for example [7, 12]) focused on other dominant fruit fly species including *Dacus bivittatus* (Bigot), *Dacus ciliatus* Loew, *Dacus punctatifrons* Karsch and *Dacus vertebratus* Bezzi. With these limited studies on fruit flies infesting cucurbit crops in Tanzania [13,15], information related to fruit fly infestations and host preferences remains limited. Yet, the extent to which fruit flies infest cucurbit crops and their host preference is vital for designing sound management strategies to suppress their population in agricultural crops Much of the necessary background information on infestation and incidences of fruit fly species infesting cucurbit crops is still lacking in Tanzanian.

Therefore, this study aimed at unravelling the infestation and incidence of fruit fly species infesting commonly grown cucurbit species (viz., watermelon, cucumber, and squash) in the plateau and mountainous zones of the Morogoro region, Tanzania. The information generated from this study is essentially important for developing strategies to suppress the population of fruit flies in Tanzania.

## 2. Materials and Methods

### 2.1. Study site

Infestation and incidence of cucurbit infesting Tephritidae (Diptera) were determined in three cucurbit crops, over three cropping seasons between March and October 2020 in Morogoro, Tanzania. The Morogoro region is located in the transition belt between S 5°58’ – S 10°0’ and E35°25’-E38°30’ [23]. The cucurbit cropping seasons included March‒April (early rain season or Season 1), June‒July (late rainy season or Season 2), and September‒October (dry season or Season 3).

Two zones in the Morogoro differing in their climatic characteristics were selected for the study. The plateau zone is located between 300-600m above sea level. Its annual rainfall ranges between 700-1200mm and an annual average temperature of 29°C. The mountainous zone is found above 600m above sea level, with an annual rainfall of 800-2500mm and a yearly temperature of ∼24°C [14,23]. Five locations, approximately 1 km apart, were identified in each agroecological zone (Table 1). Within each location for each agroecological zone, 3 plots, each measuring 1120 m2 (70 x 16 m) were established. Cucumber (*C. sativus*), variety “Ashley” was planted using a spacing of 50 x 60 cm in plot 1, watermelon (*C. lanatus*), variety ‘Sugar baby” was planted using a spacing of 1 x 1.5 m in plot 2 and squash (*C. moschata*) variety “Waltham” was planted using a spacing of 1 x 1.5 m in plot 3.

**Table 1:**
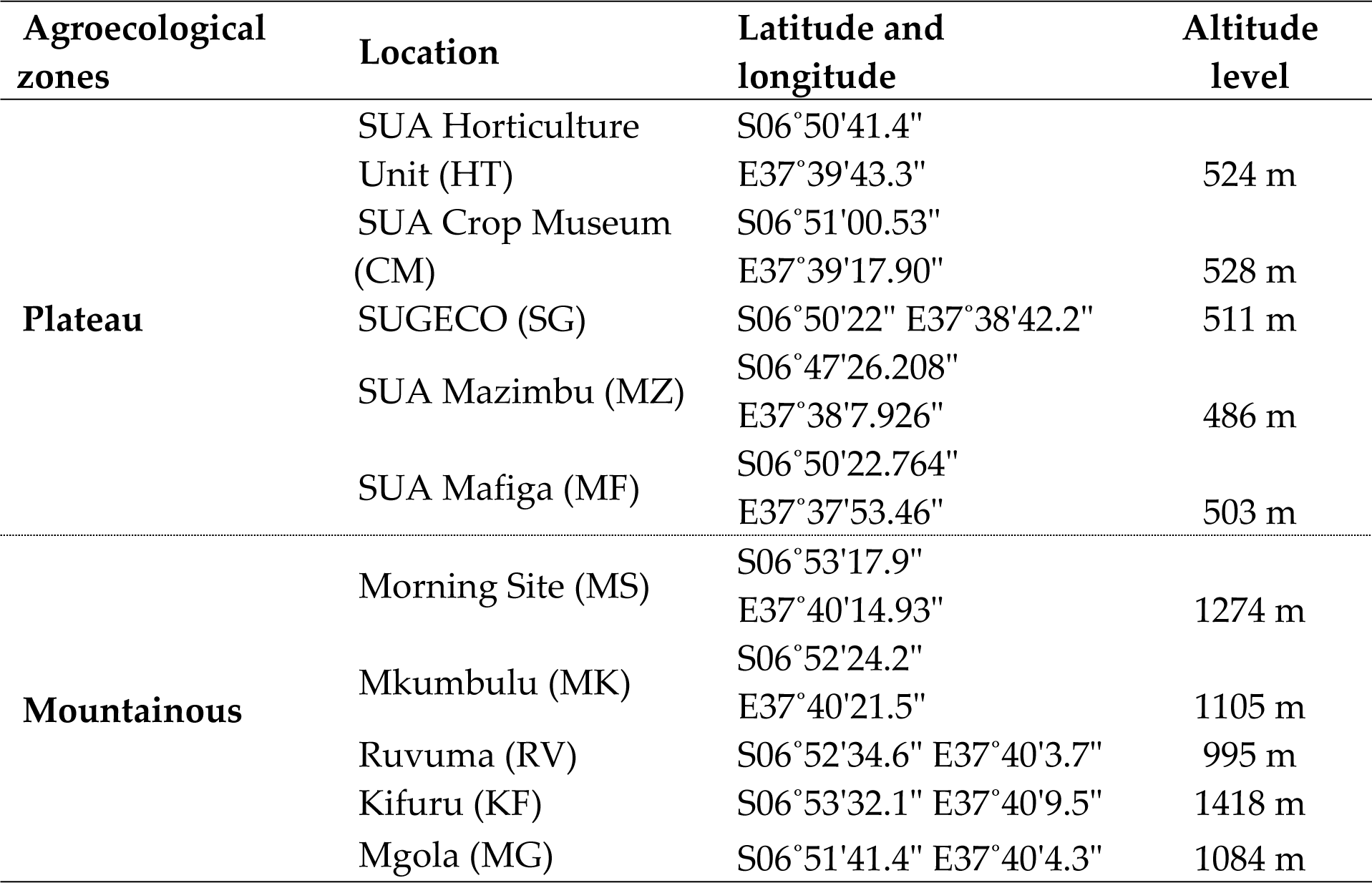
Experimental field locations in two agroecological zones of the Morogoro region.

### 2.2. Methodologies

A weekly random sampling of ten fruits (as sample unit per crop) was carried out twice per plot per week for five consecutive weeks during the peak of each fruiting season. Each fruit sample collected per cucurbit species was weighed and placed in one rearing container to rear fruit fly species infesting the collected fruits as described by [2]. The sampling was repeated across three seasons between March to October 2020. The rearing of fruit flies from infested fruits was carried out at the rearing facility of the Sokoine University of Agriculture (SUA) Morogoro using rearing containers. Containers were examined daily for adult fruit flies until no more fruit flies emerged. Room temperature ranged between 23-25°C. Emerged fruit flies were preserved in absolute ethanol for morphological identification.

### 2.3. Data collection and identification

Emerged fruit fly species per fruit sample were counted and then averaged over a week. The infestation was determined as the number of emerged adult fruit flies per kilogram (kg) per week to obtain a mean infestation per cucurbit species per week. Incidence was calculated as the number of positive samples per crop species per total number of samples times 100. It is expressed as the percentage of positive samples. Fruit flies were identified using identification keys [25, 26]. Some specimens were sent to the Royal Museum for Central Africa (RMCA) for identification.

### 2.4. Data analysis

The effects of crop species, agroecological zone and cropping season on infestations of dominant cucurbit infesters were determined using analysis of variance (ANOVA) following the general ANOVA Design (GAD) as described by [22] using the GAD package. Cochran’s C tests were used to test for homogeneity of variances [22]. The Student-Newman-Keuls (SNK) test was used for post hoc comparisons of means. All statistical analyses were performed using R -statistical software version 4.1.1 [21].

## 3. Results

### 3.1. Infestation of fruit flies among cucurbit species

Results from Table 2 show the number of positive samples per crop species for each fruit fly species. More than 85% of all cucurbit fruits were positive. In total 22 169 fruit flies were reared from the three crops. Of these 12 390 flies emerged from fruits collected in the mountainous zone, while 9 779 flies emerged from fruits collected in the plateau zone (Table 3). *Zeugodacus cucurbitae* infested the highest number of fruits. *Dacus vertebratus* and *D. bivittatus* were mostly recovered from fruits of the plateau zone while *D. punctatifrons* was only recovered from C. moschata fruits from the plateau zone. Generally, the highest incidence and infestation rates were caused by *Z. cucurbitae* followed by *D. vertebratus, D. ciliatus, D. bivittatus,* and *D. punctatifrons* (Table 3).

**Table 2:**
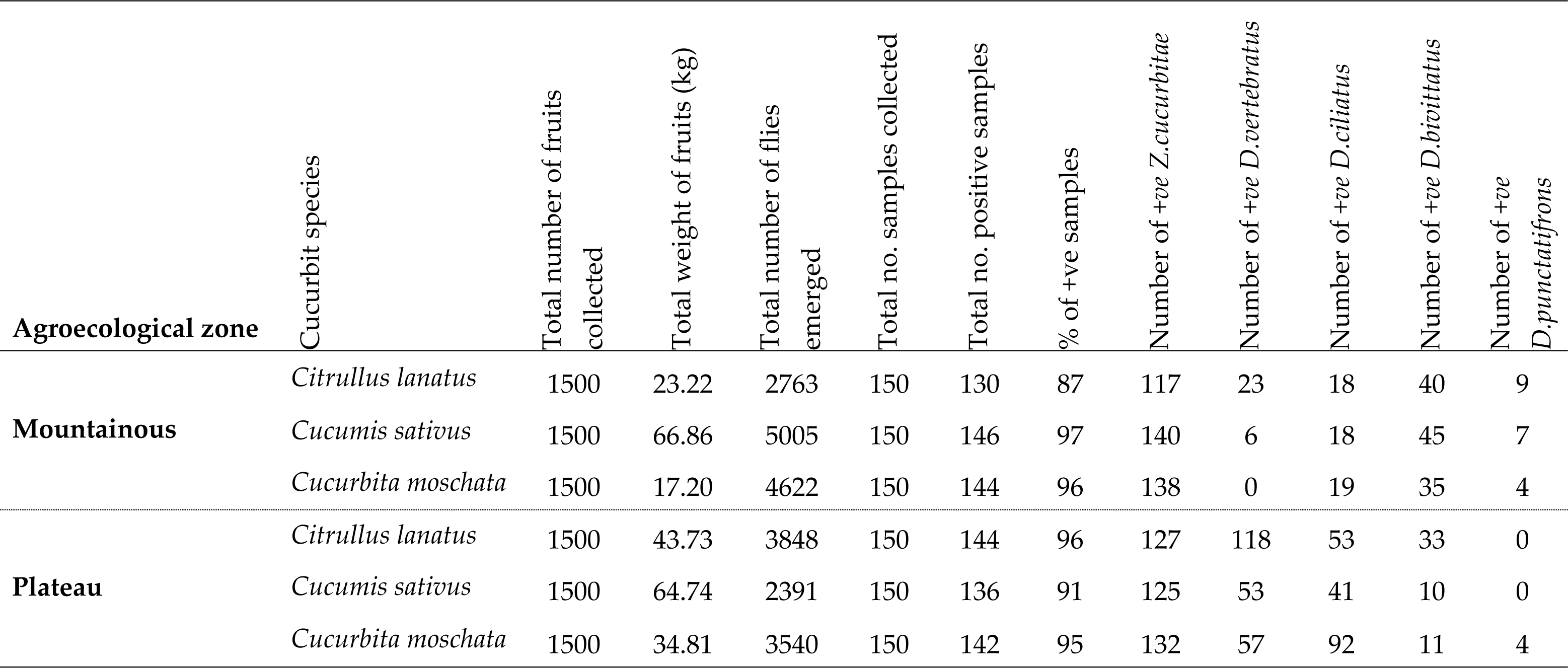
List of cucurbit species indicating positive samples for the emerged fruit fly species.

**Table 3:**
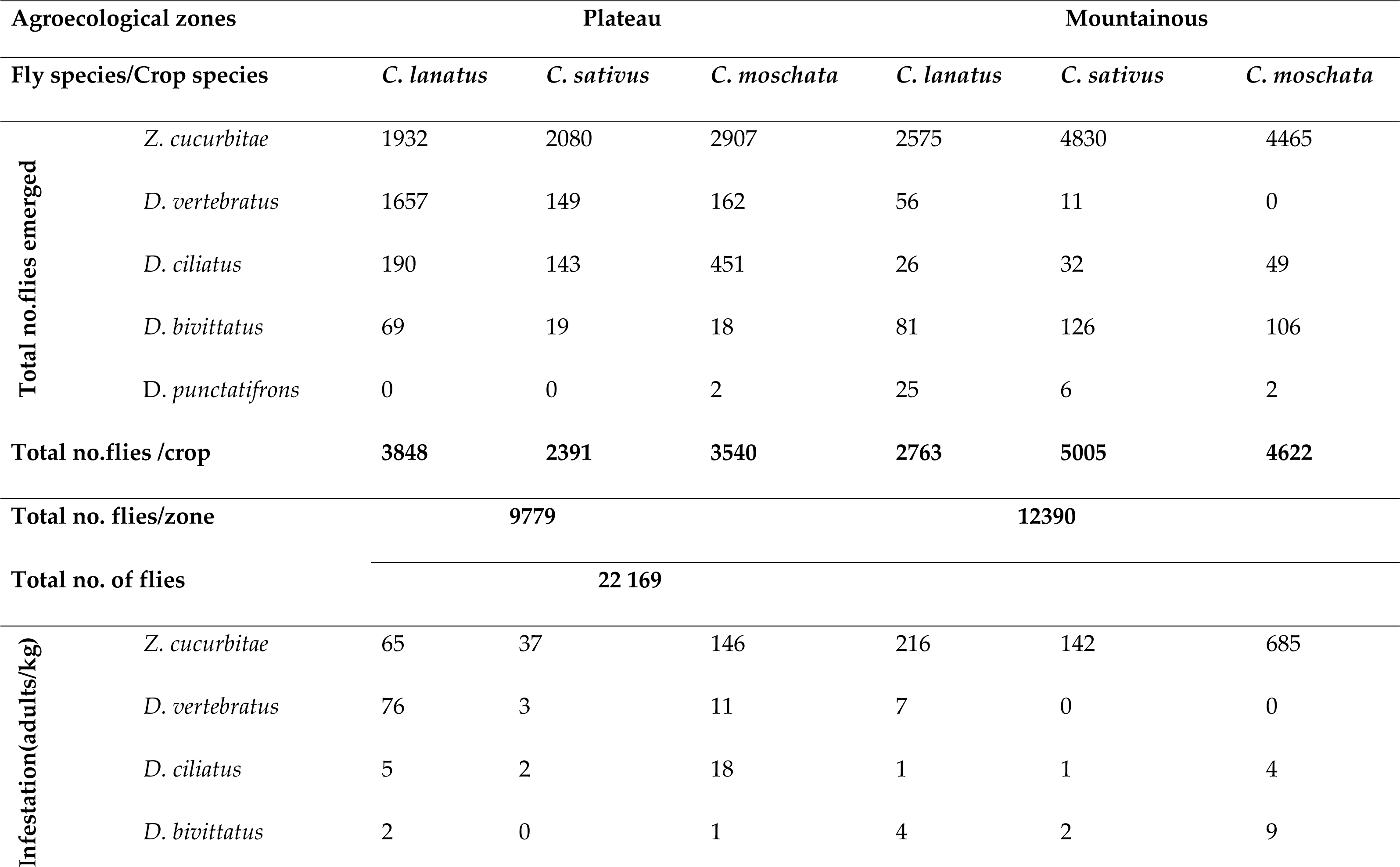
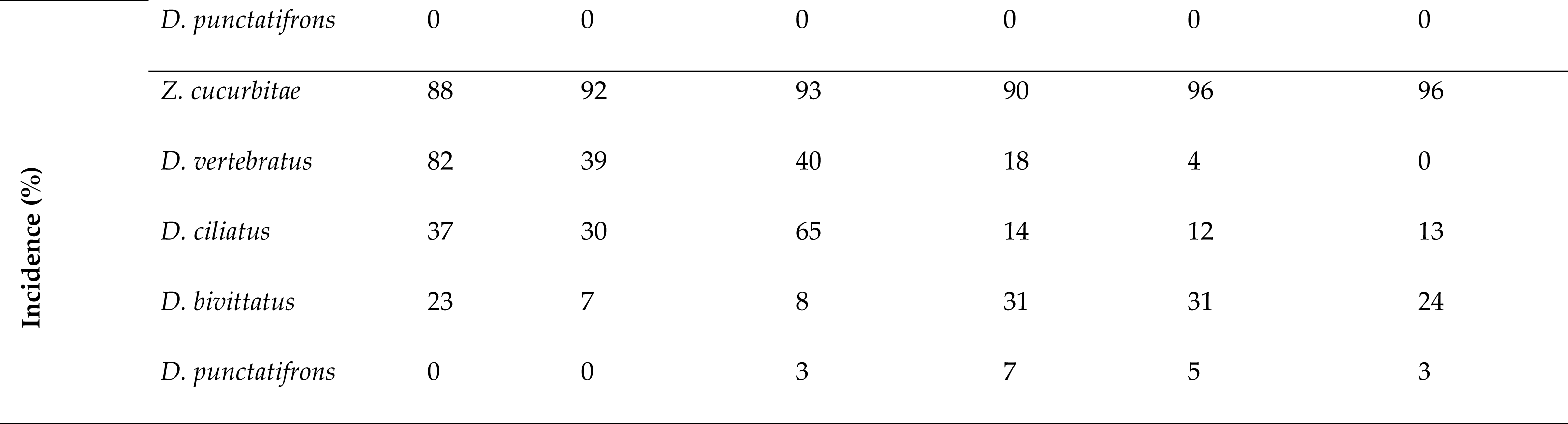
Incidence, infestation and the total number of flies reared among three cucurbit species.

### 3.2. Effect of agroecological zones, cropping seasons, and cucurbit species on the infestation of Z. cucurbitae

Results from Table 4 showed significant effects of agroecological zone × cropping season × cucurbit species on the infestation of *Z. cucurbitae* (*p* < 0.05). The *Z. cucurbitae* infestation was significantly higher in *C. moschata* than in *C. lanatus* and *C. sativus* in all agroecological zones and seasons (with only one exception) (*Post hoc* test = SNK) (Fig 1). The level of infestation was higher in *C. moschata* and *lower* in *C. sativus* across all zones and seasons.

**Fig 1.**
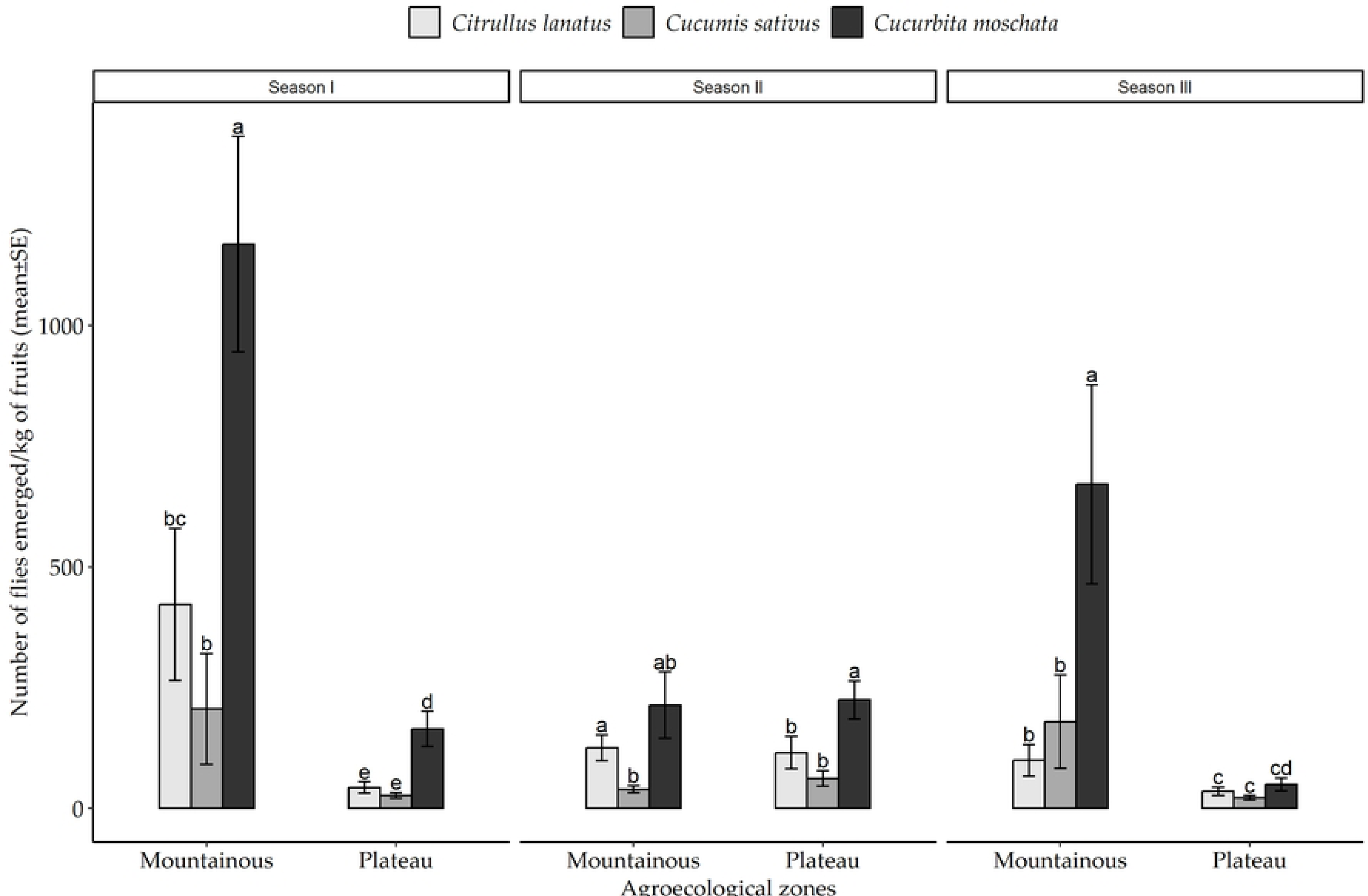
Mean infestation of *Z. cucurbitae* in different cucurbit species cultivated across the two agroecological zones of the Morogoro region from March to October 2020. Bars with different letters denote significant differences, p < 0.05. SE stands for Standard Error.

**Table 4.**
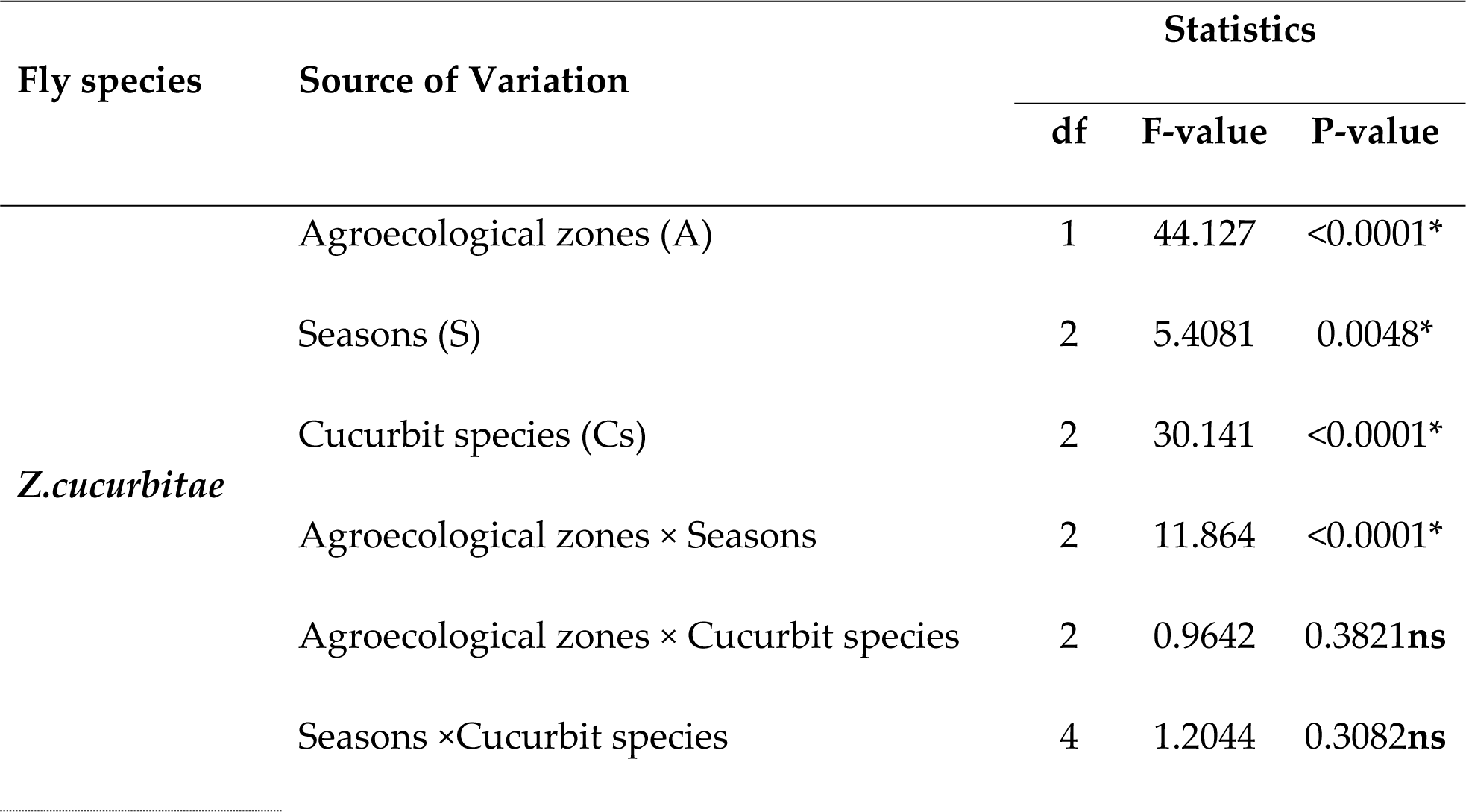
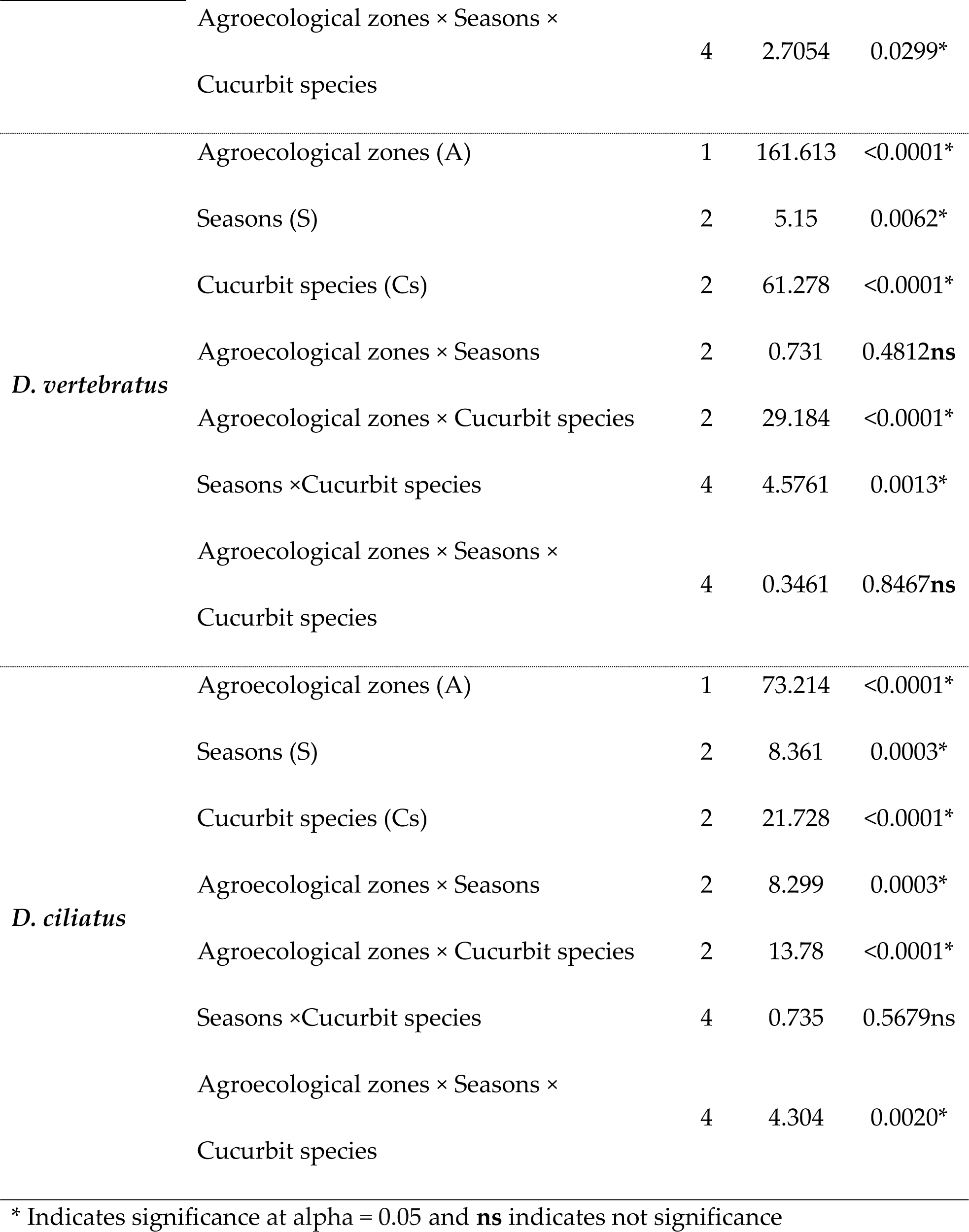
Analysis of variance for the effects of agroecological zones, cropping seasons and cucurbit species on the infestation of *Zeugodacus cucurbitae, Dacus ciliatus* and *Dacus vertebratus*.

### 3.3. Effect of agroecological zones, cropping seasons and cucurbit species on the infestation of Dacus vertebratus

Table 4 indicates significant two-way effects of agroecological zone × cucurbit species and cropping season × cucurbit species interactions on the infestation of *D. vertebratus* among cucurbit species (*p* < 0.05). The *D. vertebratus* infestation was significantly higher in *C. lanatus* than in the other two cucurbit species regardless of the season and agroecological zone (Fig 2).

**Figs 2.**
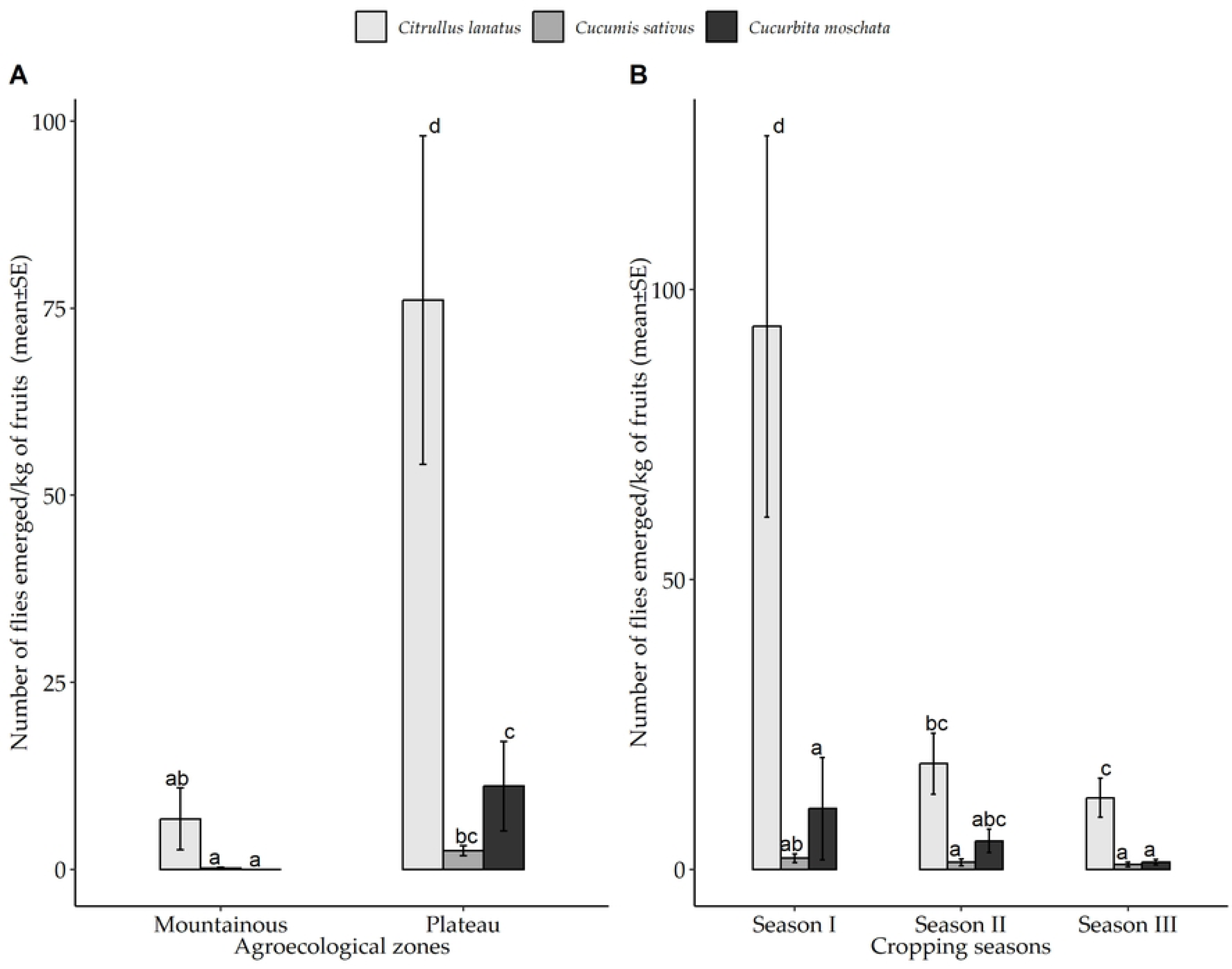
Mean infestation of *D. vertebratus* in different cucurbit species in the Morogoro. A: represents the mean infestation across the two agroecological zones and B represents the mean infestation across three cropping seasons in Morogoro from March to October 2020. Bars with different letters denote significant differences, p < 0.05. SE stands for Standard Error.

### 3.4. Effect of agroecological zones, cropping seasons, and cucurbit species on the infestation of Dacus ciliatus

Results from Table 4 showed agroecological zone × cropping season × cucurbit species interaction had significant effects on the infestation of *D. ciliatus* (*p* = 0.002). The infestation was significantly higher in *C. moschata* than in the other two cucurbit species in all seasons and across all agroecological zones (Fig 3). Generally, *C. moschata* was highly infested by *D. ciliatus* followed by *C. lanatus and C. sativus*.

**Fig 3.**
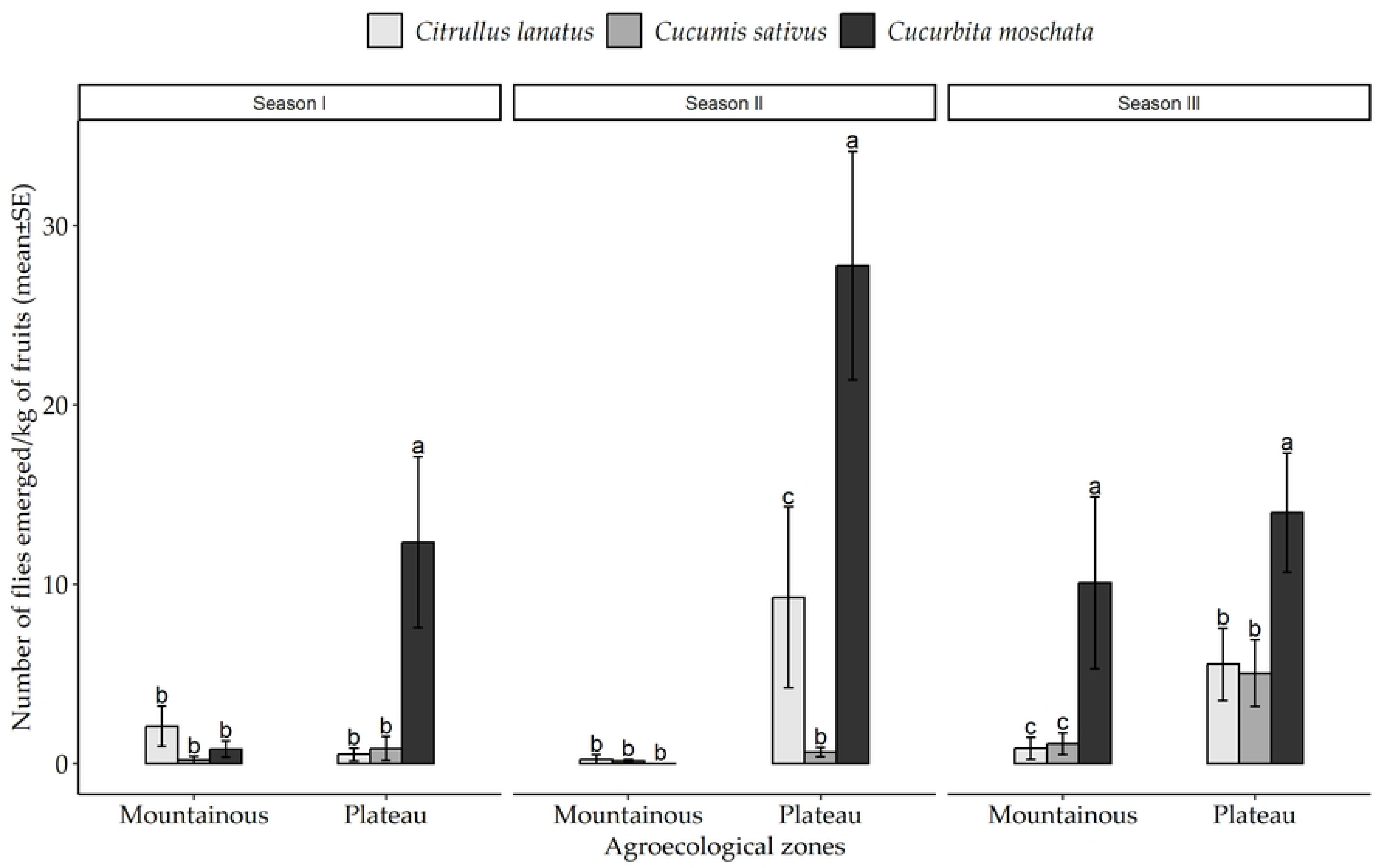
Mean infestation rates of *D. ciliatus* in different cucurbit species cultivated. across the two agroecological zones of the Morogoro region from March to October 2020. Bars with different letters denote significant differences, p < 0.05. SE stands for Standard Error.

### 3.5 Temporal and Spatial Variability of fruit fly infestation

Results indicated that all dominant flies were present in all zones, at all times over the growing seasons (Figs 4-6). We observed a large variability in infestation among the dominant fruit fly species in the study area (Figs 4-6).

**Fig 4.**
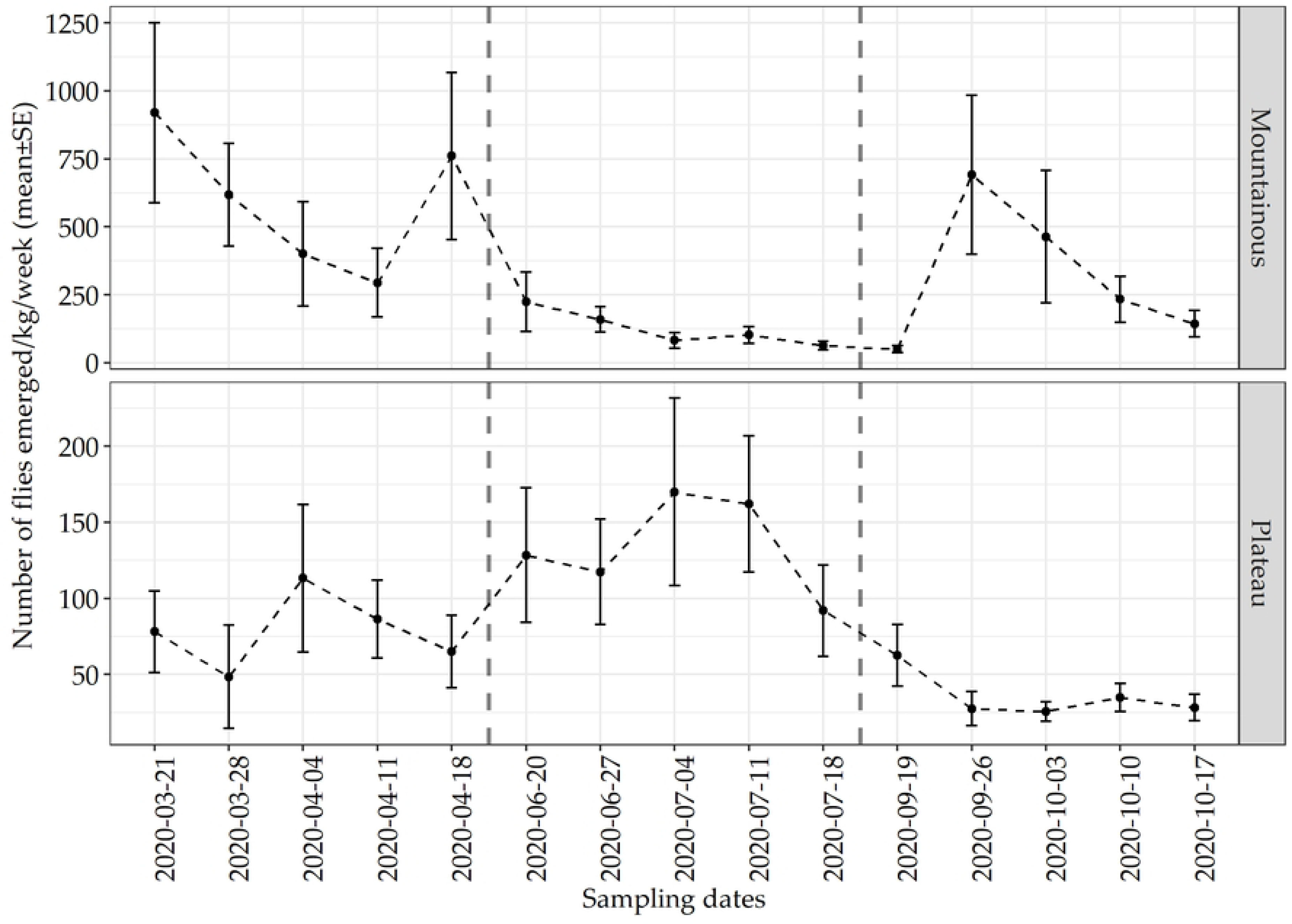
Temporal variability in the number of *Z. cucurbitae* that emerged per kilogram of fruit samples collected across the two zones of the Morogoro region from March to October 2020. The dashed lines separate seasons. SE stands for Standard Error.

*Zeugodacus cucurbitae* infestation was highest in the mountain zone in Season I (2020-03-21 to 2020-04-18), then low during Season II (2020-06-20 to 2020-07-18) and then increases in Season III (2020-09-19 to 2020-10-17) (Fig 4). For the plateau zone, the infestation was highest during season II and low in season III (Fig 4).

In the case of *D. vertebratus,* the infestation was highest in Season I, then showed low numbers in Seasons II and III across all zones (Fig 5). For *D. ciliatus,* the infestation seems to increase from Season I to II, then decrease in Season III in the plateau zone but showed very low numbers in Season I and II in the mountainous zone, which then increased in Season III.

**Fig 5.**
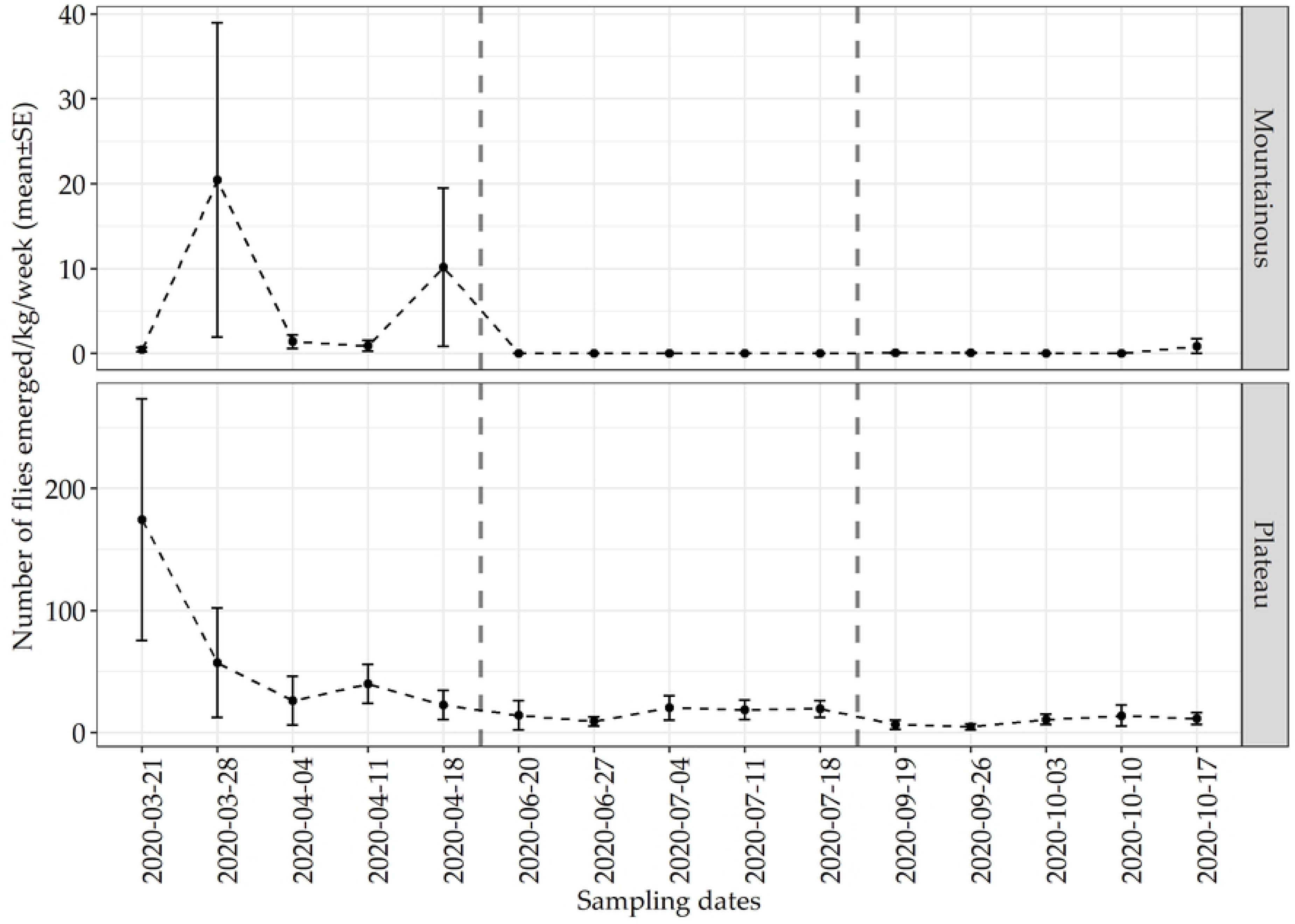
Temporal variability in the number of *D. vertebratus* emerged per kilogram of fruit samples collected across the two zones of the Morogoro region from March to October 2020. The dashed lines separate seasons. SE stands for Standard Error.

**Fig 6.**
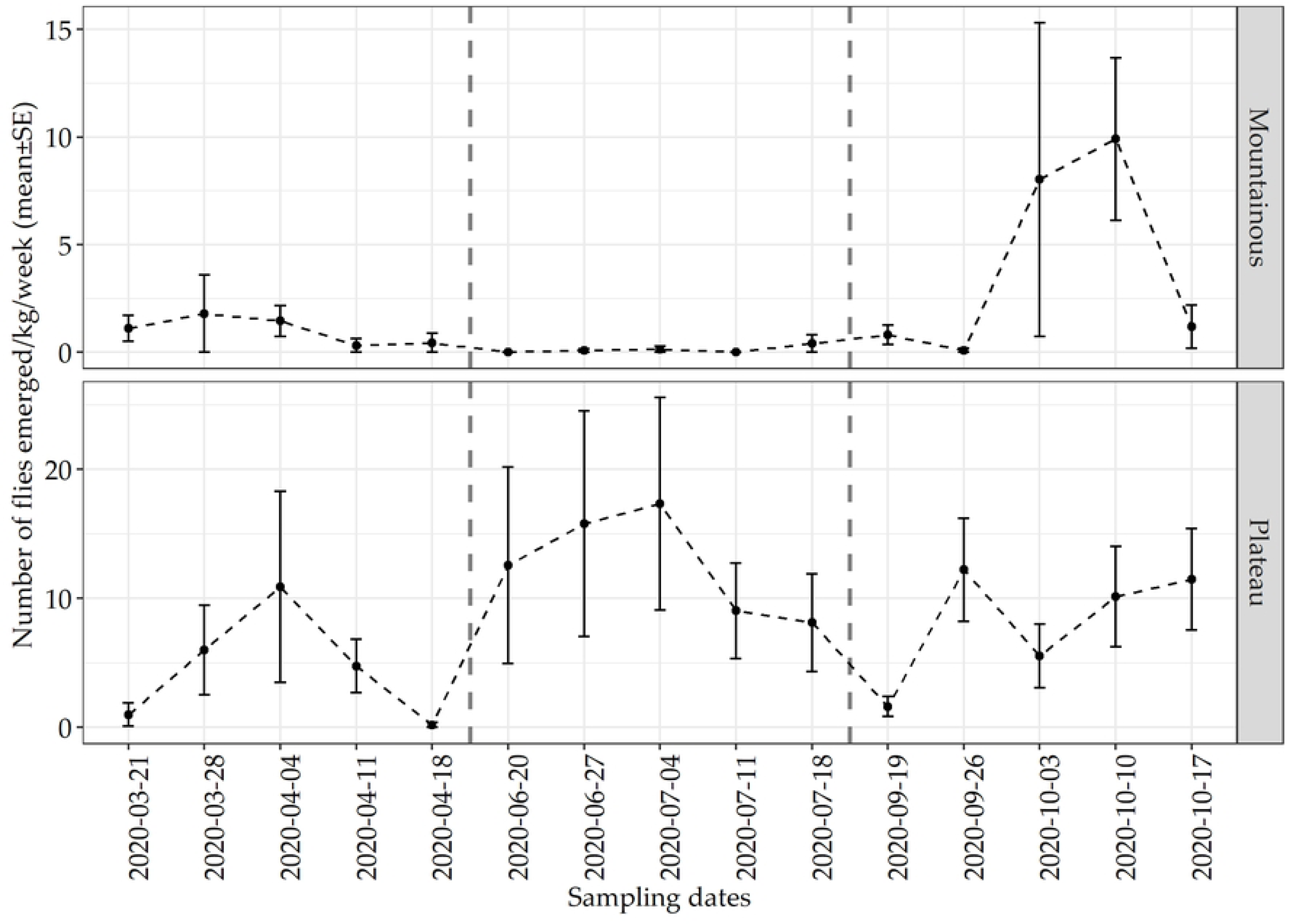
Temporal variability in the number of *D. cilliatus* emerged per kilogram of fruit samples collected across the two zones of the Morogoro region from March to October 2020. The dashed lines separate seasons. SE stands for Standard Error.

## 4. Discussion

### 4.1. Infestation and its variability

Results of this study showed that *Z. cucurbitae, D. ciliatus*, *D. vertebratus* and, to a lesser extent, *D. bivittatus* and *D. punctatifrons* are the major tephritid pests of *C. lanatus*, *C. sativus* and *C. moschata* in the Morogoro Region in Tanzania. *Zeugodacus cucurbitae* was the dominant species followed by *D. vertebratus* and *D. ciliatus*. Previous studies in East Africa by [15] in Eastern Central Tanzania and [7] in the Coastal Region of Kenya reported the dominance of *Z. cucurbitae* over native species. Similar results were reported from Benin in West Africa [6]. Infestation of *Z. cucurbitae* is however highly variable depending on the cucurbit host, seasons, and localities.

Our study indicated that Z. *cucurbitae* infestation was highest in the mountain zone in seasons I and III, and lowest in season II. In the plateau, the infestation increased from Season I to Season II and decreased in Season III. The infestation of *Z. cucurbitae* was as high as 685 flies/kg. A study by [7] reported infestation of *Z. cucurbitae* in cucurbit crops that varied from 94.1flies/kg to 624 flies/kg depending on the cucurbit species. Other studies in India reported an infestation of Z. cucurbitae of up to 431.9/kg [10]. Fruit flies normally attack immature cucurbit fruits, which are thin-skinned, small, and low-weight [12], which increases infestations.

Our study also found that *C. moschata* was the most infested species by *Z. cucurbitae* among the three cucurbit species across all agroecological zones. The overall prevalence of *Z. cucurbitae* infestations in the three species of cucurbits varied seasonally across the two agroecological zones. Such variability in infestation has been linked to host availability and climatic conditions [7,12, 13]. Our study was however limited to three commercial cucurbits and excluded many previously reported hosts [5, 7,8,12, 11, 18-20].

The present results further showed the highest infestation of *D. vertebratus* for *C. lanatus* than the other two crops at both zones and throughout the cropping seasons. A higher infestation of *D. vertebratus* for watermelon has been previously reported in other regions (see for example [7]. A study by [11] reported a high infestation of *D. vertebratus* in watermelon, and its dominance over *D. ciliatus* and *Z. cucurbitae* in the crop, which suggested interspecific competition. Our study showed large variability of *Dacus vertebratus* infestation over seasons. The highest infestation was in season I and the lowest was in seasons II and III in all zones.

*Dacus ciliatus* showed the highest infestation in C. *moschata* at both zones and between cropping seasons except in a few situations. *Citrullus lanatus* was the second most infested cucurbit species by *D. ciliatus.* A study by [7] showed that cucumber and butternut had the highest infestation of *D. ciliatus* than other cucurbit species. However, there are variable reports on the infestation of *D. ciliatus* from across regions, that nevertheless included *C. moschata* among preferred hosts as well as cucumber and watermelon [4,24].

The variability of *D. ciliatus* infestation across seasons and zones was higher. The infestation increased from seasons I to II and decreased in season III for the plateau zone but it was very low in seasons I and II in the mountainous zone, which then increased in Season III.

### 4.2. Incidence of fruit flies

Results of the present study showed up to 96% incidence of *Z. cucurbitae* in cucurbit crops. Previous studies reported a 30% to 100% incidence of *Z. cucurbitae* depending on the cucurbit species and environmental conditions [7, 14]. On other hand, [16] reported a 19.4 - 22.1% incidence of *Z. cucurbitae* in watermelons in Nepal. The present study reports up to 82% incidence of *D. vertebratus* in *C. lanatus* grown in the plateau zone, a low-altitude area of the Morogoro region. [14] report incidence values of 0%, 7.62% and 0.45% of *D. vertebratus* in watermelon, cucumber, and butternut, respectively. These figures were lower than those found in the present studies. Our results showed up to 65% incidence of *D. ciliatus* in *C. moschata*. [7] found over 70% incidence of *D. ciliatus* in squash and cucumber. Results in other crops by [10] in India reported incidences of *D. ciliatus* of up to 73.83 % in cucumber and up to 63.31 % in pickling cucumber.

## 5. Conclusion

The three dominant fruit fly species; *Z. cucurbitae*, *D. vertebratus*, and *D. ciliatus* had significantly different infestations among commonly grown cucurbit species in Morogoro. The intensity of infestation and incidence level is influenced by cucurbit species, agroecological zone and cropping season. Among the cucurbit crops, squash was the most infested by *Z. cucurbitae* and *D. ciliatus*, and watermelon by D. *vertebratus* while cucumber was the least infested species by all dominant fruit fly species.

In general, this study confirms the destructive potential of *Z. cucurbitae, D. vertebratus*, and *D. ciliatus* toward the commonly grown cucurbit crops in Morogoro, and addresses the need to consider the variability of infestation over seasons and localities in developing measures to reduce their current populations.

## Acknowledgements

We thank Cessila Kijalo, Evance Matowo, Frida Mangu, Christian Barnabas, and Tabitha Fussi (Sokoine University of Agriculture) for their assistance in the field and the lab, to I.M. White (Natural History Museum UK) for making available his forthcoming revision on dacines of the Afrotropical region. We also thank Wouter Hendrycks and Lore Esselens for their help in the analysis of the data.

## References

1. Am, M., Sridharan, C. S., Awasthi, N. S. Varying infestation of fruit fly, *Bactrocera cucurbitae* (Coquillett) in different cucurbit crops. Journal of Entomology and Zoology Studies 2017, 5(53), 1419–1421.

2. Copeland, R.S., Wharton, R.A., Luke, Q. De Meyer, M. Indigenous hosts of *Ceratitis capitata* in Kenya. Annals of Entomological Society of America 2002, 95, 672–694.

3. Deguine, J. P., Atiama-Nurbel, T., Aubertot, J. N., Augusseau, X., Atiama, M., Jacquot, M. Reynaud, B. Agroecological management of cucurbit-infesting fruit fly: a review. Agronomy for Sustainable Development, 2015, 35, 937–965. DOI: https://doi.org/10.1007/s13593-015-0290-5

4. Deguine, J. P., Atiama-Nurbel, T., Douraguia, E., Chiroleu, F., Quilici, S. Species diversity within a community of the cucurbit fruit flies *Bactrocera cucurbitae*, Dacus ciliatus, and Dacus demmerezi roosting in corn borders near cucurbit production areas of Reunion Island. Journal of Insect Science 2012, 12,1–32. DOI; https://doi.org/10.1673/031.012.3201

5. Dhillon, M.K., Singh, R., Naresh, J.S. and Sharma, H.C. The melon fruit fly, Bactrocera cucurbitae: A review of its biology and management. Journal of Insect Science 2005, 5, 40 – 57. DOI: https://doi.org/10.1093/jis/5.1.40

6. Gnanvossou, D., Hanna, R., Goergen, G., Salifu, D., Tanga, C. M., Mohamed, S. A., Ekesi, S. Diversity and seasonal abundance of tephritid fruit flies in three agro-ecosystems in Benin, West Africa. Journal of Applied Entomology 2017, 141(10), 798–809. DOI:https://doi.org/10.1111/jen.12429

7. Kambura, C., Tanga, C.M., Kilalo, D., Muthomi, J., Salifu, D., Rwomushana, I., Mohamed, S.A., Ekesi, S. Composition, Host Range and Host Suitability of Vegetable-Infesting Tephritids on Cucurbits Cultivated in Kenya. African Entomology 2018, 26(2), 379–397.

8. Layodé, B. F. R., Onzo, A., Karlsson, M. F. Watermelon-infesting Tephritidae fruit fly guild and parasitism by Psyttalia phaeostigma (Hymenoptera: Braconidae). International Journal of Tropical Insect Science 2020, 40,157–166. DOI: https://doi.org/10.1007/s42690-019-00066-x

9. Loukou, A. L., Gnakri, D., Djè, Y., Kippré, A. V., Malice, M. J. P. B., Baudoin, J. P., Bi, I. A. Macronutrient composition of three cucurbit species cultivated for seed consumption in Côte d’Ivoire. African Journal of Biotechnology 2007, 6(5),1–5.

10. Manoj, A., Sridharan, S., Mohan, C., Nikita, S.A. Varying infestation of fruit fly, *Bactrocera Cucurbitae* (Coquillett) in Different Cucurbit Crops. Journal of Entomology and Zoology Studies 2017, 5(3),1419–1421.

11. Materu, C. L., Losujaki, E. W., Zain, I. Farmers Knowledge on Integrated Pest Management in Cucurbit Production. International Journal of Research-GRANTHAALAYAH 2018, 6(12), 70–76.

12. Mokam, D. G., Bilong Bilong, C. F., Djiéto-Lordon, C., Lumaret, J. P. Host susceptibility and pest status of fruit flies (Diptera: Tephritidae) attacking cucurbits in two agroecological zones of Cameroon, Central Africa. African Entomology 2018, 26(2),317–332. DOI: doi/epdf/10.4001/003.026.0317

13. Mwatawala M., Maerere, A., Makundi R.H. De Meyer, M. Seasonality and host plant preference of *Bactrocera cucurbitae* in Central Tanzania. International Journal of Pest Management 2010, 56(3), 256–273. DOI; https://doi.org/10.1080/09670871003596792

14. Mwatawala, M.W. De Meyer, M., Makundi, R.H. Maerere, A.P. Host range and distribution of fruit-infesting pestiferous fruit flies (Diptera: tephritidae) in selected areas of Central Tanzania. Bulletin of Entomological Research 2009, 99, 629–641. DOI: https://doi.org/10.1017/S0007485309006695

15. Mwatawala, M.W., Abdul, K., Godfrey, E., Jeremiah, S., Virgilio, M. De Meyer, M. Preference of *Bactrocera cucurbitae* Coquillett for three commercial fruit vegetable hosts in natural and semi-natural conditions. Fruits 2015, 70 (6), 333 – 339. DOI: https://doi.org/10.1051/fruits/2015034

16. Pradhan, R.B. Relative susceptibilities of some vegetables grown in Kathmandu valley to *D. cucurbitae* Coq. Nepal Journal of Agriculture 1976, 12, 67–75.

17. Ryckewaert, P., Deguine, J.P., Brévault, T. Vayssières, J.F. Fruit flies (Diptera: Tephritidae) on vegetable crops in Reunion Island (Indian Ocean): state of knowledge, control methods and prospects for management. Fruits, 2010, 65(2), 113–130. DOI: https://doi.org/10.1051/fruits/20010006

18. Sapkota, R. Damage assessment and field management of cucurbit fruit fly (*Bactrocera cucurbitae* Coquillett) in squash during the spring-summer season of mid-hill Nepal. Thesis, M. Sc. Ag., Tribhuvan University/IAAS, Rampur, Nepal; 2009.

19. Shooker, P., Khayrattee, F., Permalloo, S. Use of maize as a trap crop for the control of melon fly, *Bactrocera cucurbitae* (Diptera: Tephritidae) with GF-120. Bio-control and other control methods (online); 2006. Available on: http://www.fela.edu/flaEnt/fe87p3s4.pdf.

20. Sohrab, W. H., Prasad, C. S. Investigation on level of infestation and management of cucurbit fruit fly, Bactrocera cucurbitae (Coquillett) in different cucurbit crops. International Journal of Pure Applied Bioscience SPI 2018, 6(1), 184–196.

21. Team, R. C. R. A language and environment for statistical computing. Published online 2020, 2021.

22. Underwood, A.J. *Experiments in Ecology: Their Logical Design and Interpretation Using Analysis of Variance.* Cambridge University Press, Cambridge; 1997. DOI: https://doi.org/10.1016/S0022-0981(00)00181-7

23. United Republic of Tanzania (URT). Morogoro Regional Socio-Economic Profile. National Bureau of Statistics Morogoro Regional Commissioner’s office, Morogoro; 2002, 229pp.

24. Vayssières, J. F., Carel, Y., Coubès, M., Duyck, P. F. Development of immature stages and comparative demography of two cucurbit-attacking fruit flies in Reunion Island: *Bactrocera cucurbitae* and *Dacus ciliatus* (Diptera Tephritidae). Environmental Entomology 2008, 37(2), 307–314.

25. Virgilio M, White IM, De Meyer M. A set of multi-entry identification keys to African frugivorous flies (Diptera, Tephritidae). ZooKeys 2014, 428,97–108. DOI: https://doi.org/10.3897/zookeys.428.7366

26. White, I.M. Taxonomy of the Dacina (Diptera: Tephritidae) of African and the Middle East. African Entomology Memoir 2006,2, 1–156.

